# A natural language processing system for the efficient extraction of cell markers

**DOI:** 10.1101/2024.01.30.578115

**Authors:** Peng Cheng, Yan Peng, Xiao-Ling Zhang, Sheng Chen, Bin-Bin Fang, Yan-Ze Li, Yi-Min Sun

## Abstract

1.

**Background:** In the last few years, single-cell RNA sequencing (scRNA-seq) has been widely used in various species and tissues. The construction of the cellular landscape for a given species or tissue requires precise annotation of cell types, which relies on the quality and completeness of existing empirical knowledge or manually curated cell marker databases. The natural language processing (NLP) technique is a potent tool in text mining that enables the rapid extraction of entities of interest and relationships between them by parsing the syntax structure.

**Methods and results:** We developed MarkerGeneBERT, an NLP-based system designed to automatically extract information about species, tissues, cell types and cell marker genes by parsing the full texts of the literature from single-cell sequencing studies. As a result, 8873 cell markers of 1733 cell types in 435 human tissues/subtissues and 9064 cell markers of 1832 cell types in 492 mouse tissues/subtissues were collected from 3987 single-cell sequencing-related studies. By comparison with the marker genes of existing manual curated cell marker databases, our method achieved 76% completeness and 75% accuracy. Furthermore, within the same literature, we found 89 cell types and 183 marker genes for which the cell marker database was not available. Finally, we annotated brain tissue single-cell sequencing data directly using the compiled list of brain tissue marker genes from our software, and the results were consistent with those of the original studies. Taken together, the results of this study illustrate for the first time how systematic application of NLP-based methods could expedite and enhance the annotation and interpretation of scRNA-seq data.

## 2. Introduction

Single-cell sequencing technology, which has exceptional singular cell resolution, has pioneered a burgeoning field of research across numerous species and tissues[1]. The primary advantage resides in its capacity to precisely outline the cellular landscape—comprehensive maps of all cell types within distinct tissues and organs. For comprehensive annotation of various cell types, researchers must first identify the potential cell types present in the tissue and then aggregate the corresponding cell type marker genes through comprehensive literature reviews or by referencing existing cell marker databases. Additionally, tools such as CellAssign[2] and scCATCH[3], which are mechanical tools for coarse-grained annotation, are also based on existing cell marker databases or custom-built databases[4–6].

Indeed, there are already multiple databases encompassing cell markers for various species and tissue types, such as CellMarkerV2[7], PanglaoDB[8], singleCellBase[9], PCMDB[10], and CancerSEA[11]. The authors of these databases collected cell marker genes by manually reviewing and curating conclusive sentences from the main text or figures of scientific articles, with the advantage of generally obtaining more accurate marker genes. However, this manual reading and curation approach requires considerable human effort and time.

Numerous text mining-based methodologies have been implemented in various research fields for identifying entities of interest and discerning the relationships between these entities by parsing syntactic dependencies within the text. For instance, Shetty *et al*. developed a language model called MaterialsBERT, which was trained using 2.4 million abstracts from the polymer literature to autonomously extract various properties of organic and polymer materials from the literature abstract[12]. Gu *et al*. employed a pretrained NLP text mining system called MarkerGenie to identify entities of interest, such as diseases, microbiomes, genes, and metabolites, that were mentioned in texts. After entity identification, the system parses the syntactic structure of the text and extracts contextual features for each word, thereby distinguishing between the types of relationships—diagnostic, predictive, prognostic, predisposing, or treatment related—among diseases, microbiomes, genes, and metabolites[13]. Naseri *et al*. utilized an NLP pipeline to identify pain-related medical terms from largely unstructured and non-standardized clinical consultation notes, subsequently predicting pain scores based on recognized pain terms[14]. Doddahonnaiah *et al*. utilized a precompiled cell type and gene vocabulary to assess the correlation between gene and cell type entities by calculating their co-occurrence frequency within more than 26 million biomedical documents[15]. In conclusion, these published methods provided a more efficient and comprehensive analysis of research articles than manual curation by aiding in the identification of rare or novel entities of interest along with their interrelations.

In this study, we developed an NLP-based cell marker extraction system called MarkerGeneBERT. MarkerGeneBERT leverages existing biomedical corpora such as CRAFT[16], JNLPBA[17], and BIONLP13CG[18] to automatically identify cell and gene entities. Furthermore, MarkerGeneBERT introduces a text classification model aimed at mitigating the false positive rate of pairwise associations between human protein-coding genes and cell types. This text classification model was based on the manual curation of 27323 sentences as pretrained data, which contain both cell and gene entities and were categorized according to whether they had marker genes. We collected 3987 single-cell sequencing articles from free-text PubMed and PubMed Central from 2017 to June 2023 and input them into MarkerGeneBERT to extract cell marker genes. Subsequently, we validated the comprehensiveness and accuracy of our identified cell marker genes by comparing them with results from manually curated cell marker databases. Furthermore, we applied scCATCH for cell cluster annotation in brain tissue samples based on our marker gene list, and the annotation results were consistent with those of previous studies. An overview of MarkerGeneBERT is given in Figure 1, which consists of four main components: literature retrieval, extraction of marker-related sentences, establishment of cell-marker associations, and inference of species, tissue, and disease information within the articles.

**Fig. 1.**
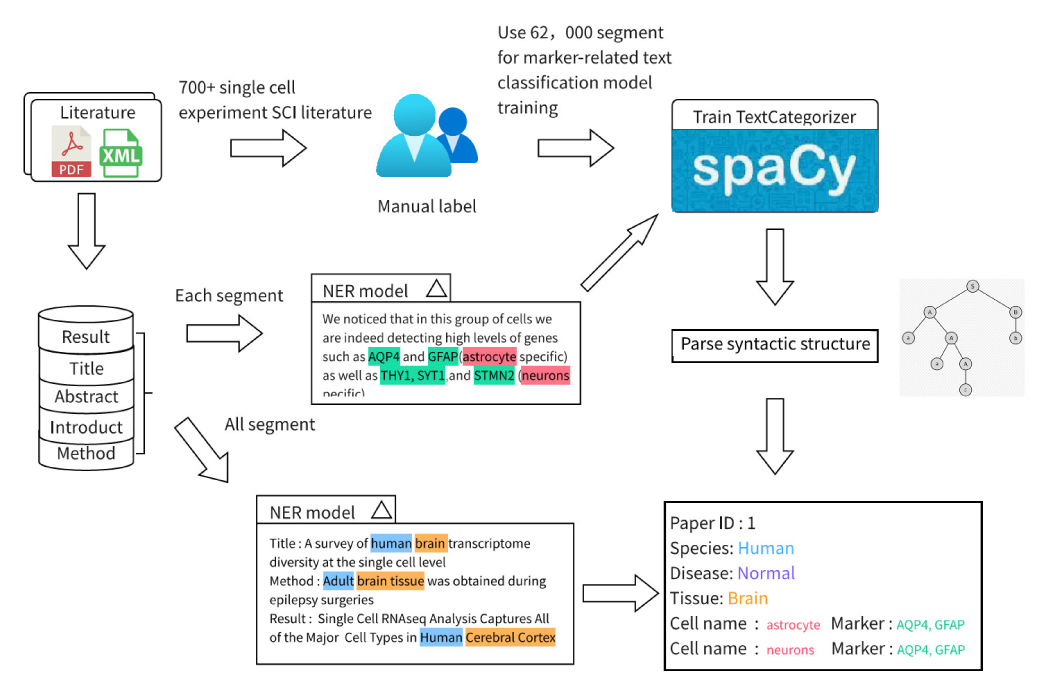
The pipeline of MarkerGeneBERT for extracting cell marker genes from the literature.

## 3. Method

### 3.1 Data collection

We systematically downloaded and parsed the main texts of approximately 20000 studies from free-text PubMed and PubMed Central. Specifically, we employed the R package “RISmed”[19] to retrieve literature using the search terms “Animals”[MeSH Terms] AND “Single-Cell Analysis”[MeSH Terms] OR “single-cell” AND “expression” within a specified time frame. These rigorous rules enabled us to obtain a comprehensive collection of PMID from single-cell research-related studies. Subsequently, using the R package “easyPubMed”[20], we acquired basic information such as titles, abstracts, and literature sources for each study. For the literature sourced from the PMC, we utilized the R package “europepmc” to retrieve the main text documents and systematically extracted sections, including the introduction, method, and results. For other manually collected literature in PDF format, we employed the python library “scipdf_parser” to parse the PDF files and extract pertinent sections such as the introduction, method, and results based on the parsed outcomes.

### 3.2 Marker-related sentence classification model

#### 3.2.1 Supervised training data generation for marker-related sentence classification

To identify marker-related sentences in the main text of the literature, specifically concerning those containing both cell and gene names with a particular syntactic structure, such as “Gene A is a marker of Cell B” or “Gene A (specific to Cell B)”, we constructed a text classification model based on the spaCy[21] and “textcat” modes pretrained on a manually annotated marker-related dataset curated by our team.

A total of 62,000 main text sentences were initially collected from free-text in PubMed and PubMed Central to generate the training data. If any of these sentences contained cell name abbreviations, they were further expanded. The expansion process was realized by inputting all the sentences from each source document into the “abbreviation_detector” component of the scispacy[22], which is capable of recognizing all abbreviations and their corresponding full forms in the text; then, the abbreviations in the sentences were replaced with their complete vocabulary.

There were more than ten bioinformatics engineers specializing in single-cell research to manually screen for cell-marker-gene-containing sentences from the raw sentences containing both cells and genes. Following the first processing step, 62000 initial sentences were narrowed down to 27323 remaining sentences. Subsequently, the 27323 sentences were randomly shuffled and reassigned to the aforementioned bioinformatics engineers utilizing predefined rules (Table 1) and their personal expertise to conduct manual labeling. The annotated sentences were then collated and reviewed by two additional senior bioinformatics engineers. Sentences with disputed annotations were discussed and subjected to re-annotation.

**Table 1.**
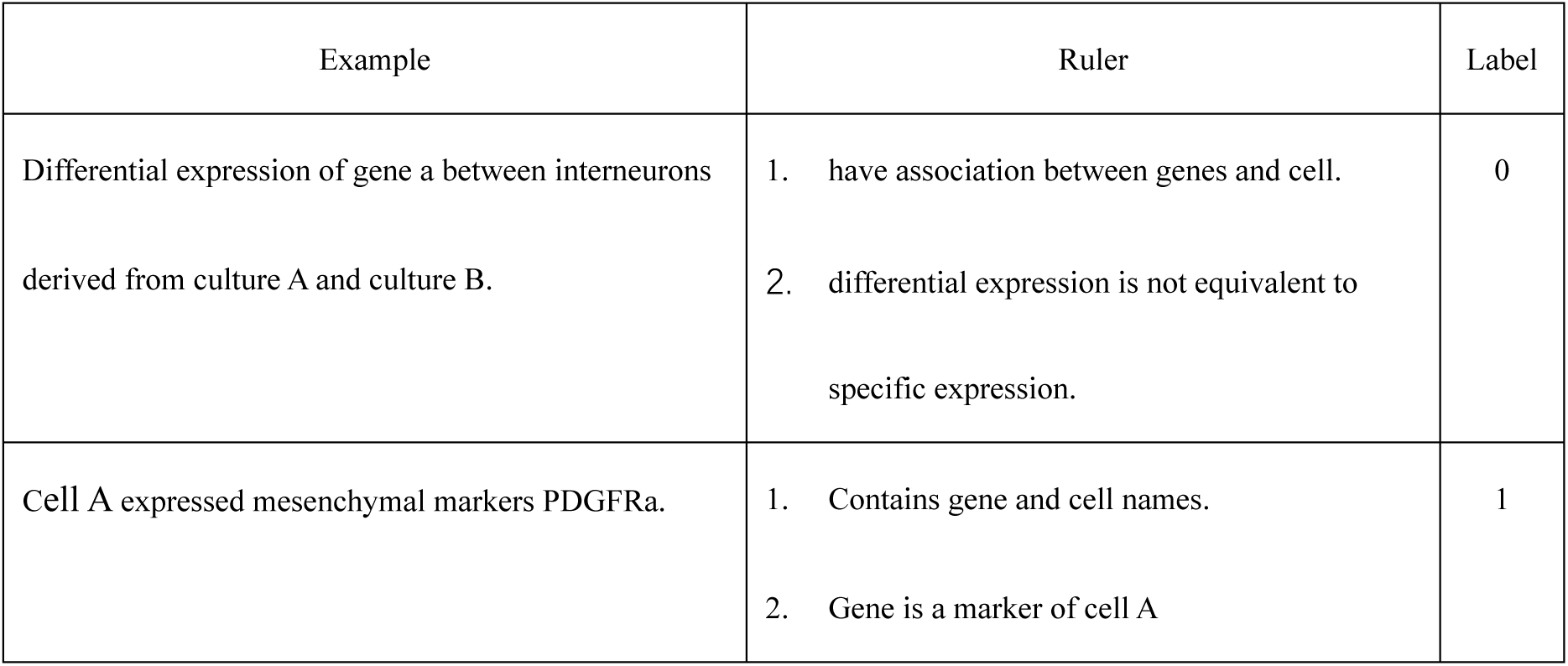
Marker-related sentence annotation rules.

#### 3.2.2 Text preprocessing of marker-related sentences

Text preprocessing has been a traditionally important step for NLP tasks. It transforms text into a more digestible form so that machine learning algorithms can perform better. Specifically, the sentences were initially input into the SciBERT model[23], and the tokenizer and parser components of the SciBERT model were used for part-of-speech tagging and syntactic dependency parsing of the sentences. This process generated contiguous spans of tokens, including words, punctuation symbols, and whitespace. Non-gene entity tokens were subsequently lemmatized and converted to lowercase characters. Additionally, tokens classified as stop words, punctuation marks (excluding parentheses), or numerical values were filtered out. Lastly, based on prior knowledge, we selectively retained parentheses only when the token inside or preceding the parentheses constituted a gene name.

We performed text preprocessing on each sentence, and the cleaned sentences were exclusively utilized for training a text classification model.

#### 3.2.3 Marker-related sentence classification model construction

The cleaned 27323 sentences were utilized as the training dataset to training a marker-related sentence classification model. The classification model was developed using spaCy’s TextCategorizer (TextCat), a powerful component within the spaCy natural language processing library, including bag-of-words model and a neural network model, characterizing authenticity through probability values in it’s output.

To determine an appropriate threshold for distinguishing marker-related sentences, the original training dataset was evenly divided into 10 parts, ensuring a 1:1 ratio of label 0 and label 1 in each subset. Subsequently, a 10-fold cross-validation approach was employed, where 9 parts of the original training set were used as a new training set to train the text classification model, while the remaining 1 part served as the validation set for evaluating the model’s performance. The sentences from the validation set were inputted into the model, yielding a predicted probability corresponding to the likelihood of the sentence being a marker-related sentence. We evaluated the precision and recall of the model at different probability thresholds and calculated the F1 score. Finally, based on the variations in F1 score under different thresholds, an appropriate threshold was selected.

### 3.3 Entity extraction

#### 3.3.1 Named Entity Recognition (NER)

To extract the entity information from the text, such as cells, species, tissues, and disease names, we imported four NER models from sciSpacy. Each NER model was originally constructed based on different biomedical corpora, enabling the recognition of distinct entity types. As shown in the Table 2, we employed different models or combinations of models to extract cell, species, tissue, and disease entities.

**Table 2.** NER model for identifying different entity types.

|  | Cell type | Species | Tissue | Disease |
| --- | --- | --- | --- | --- |
| en_ner_jnlpba_md | √ | - | - | - |
| en_ner_craft_md | √ | √ | - | - |
| en_ner_bionlp13cg_md | √ | - | √ | - |
| en_ner_bc5cdr_md | - | - | - | √ |

##### 3.3.1.1 Generation of gene vocabulary

The complete set of human and mouse protein-coding genes was obtained from the GTF file in Cell Ranger v 5.0.1, and gene entities were extracted using exact string matching.

##### 3.3.1.2 Cell entity recognition

First, each sentence was parsed, and cell names were extracted using three NER models independently (Fig. 2). Specifically, the “en_ner_craft_md” model identified entities with the entity type “CL” as cell names, the “en_ner_jnlpba_md” model recognized entities with the entity types “CELL_TYPE“ and “CELL_LINE“ as cell names, and the “en_ner_bionlp13cg_md” model identified entities with the entity type “CELL” as cell names. Subsequently, we performed exact string matching on the same sentence using the comprehensive cell names obtained from the cell ontology database[24]. Finally, we retained the cell names that were extracted by at least two sources (three models and exact string matching) as the cell names present in the respective sentences.

**Fig. 2.**
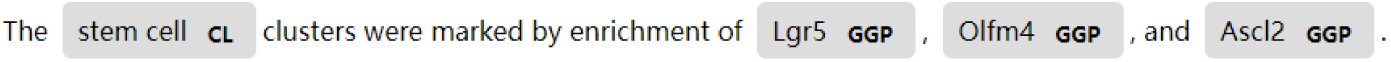
An example of an NER model for identifying and classifying named entities. CL indicates the cell line, and GGP indicates the gene or gene product.

Considering that models might not capture all the cell names comprehensively, for instance, in the case of text containing “CD4+ T cell”, some models might extract the complete cell entity as “CD4+ T cell”, while others may only extract “T cell” as the cell entity. To address this issue, we compared and completed the cell names identified by different models at the same position in the text. For example, if there were two models in which “CD4+ T cell” and “T cell” as cell entities at the same position in the text, we considered the extracted “T cell” as “CD4+ T cell”.

##### 3.3.1.3 A full-text-based strategy for extracting species and tissue entities

We employed a full-text-based strategy in which the literature was divided into sections such as abstracts, methods, and results, and entity recognition was performed using NER models on each section, followed by comprehensive analysis and judgment.

###### 3.3.1.3.1 Species entity recognition

The extraction of species entities primarily relied on MeSH (Medical Subject Headings) terms, which are controlled vocabulary thesaurus used by the National Library of Medicine (NLM) for indexing articles in PubMed. For each study, we utilized the “en_ner_craft_md” model to identify species entities from the MeSH terms. If no species entities were identified from the MeSH term text provided by PubMed, we further performed species entity recognition based on the overall structure of the full text. Specifically, we employed the “en_ner_craft_md” model to identify species entities separately from the text in the title section, methods section, and first paragraph of the Results section. The most frequently occurring species entity was selected as the species type studied in the respective literature.

###### 3.3.1.3.2 Tissue entity recognition

We utilized the “en_ner_bionlp13cg_md” model for the recognition of tissue entities. Specifically, for each study, we identified tissue entities separately from the MeSH term text, the text in the title section, and the sentences within the full text that contained keywords related to single-cell sequencing, such as “single-cell” and “dissociation”. If a tissue entity was identified in all three text sections of the literature, it was considered the correct tissue type. Otherwise, we supplemented the recognition of tissue entities by analyzing the text in the first paragraph of the Results section and the Methods section of the article. We calculated the frequency of each entity extracted from different text sources and ranked them accordingly. Additionally, we determined the frequency of co-occurrence between each entity and keywords related to single-cell sequencing in the same sentence, as well as the frequency of co-occurrence between each entity and all cell entities identified in the literature. The top two tissue types based on the cumulative ranks from these three ranking results were considered candidate tissue types according to the literature.

##### 3.3.1.4 Disease entity recognition

We employed the “en_ner_bc5cdr_md” model to identify disease entities from the title section, which were considered the disease types studied in the respective literature. If no disease entities were detected, the literature was assumed to be “normal” by default.

### 3.4 Cell type–gene relation classification

To obtain cell marker genes from marker-related sentences, we first retained sentences that simultaneously contained cell and gene names based on the results of entity recognition. Next, the sentences were subjected to text cleaning and input into a text classification model. If the predicted probability value exceeded a certain threshold, the original sentences were further categorized into two types: those suitable for extracting cell–gene relationship pairs based on rules and those requiring manual extraction of cell–gene relationship pairs (Table 3).

**Table 3.** Classification of sentence extraction methods.

| relation classification | Keyword | Example |
| --- | --- | --- |
| Suitable |  | We noticed that in this group of cells we are indeed detecting high levels of genes such as AQP4 and GFAP (astrocyte specific) as well as THY1, SYT1, and STMN2 (neuron specific) |
| Manual extraction required | Small, non, not, non, respectively, neither, Neither, distinguish, nor, no, decreased, decrease, downregulated, downregulate, weak, absent, absented, low, lack, lacks, lower, few | The differentiated subcutaneous adipocytes from Fam13a KO mice also expressed a marginally but not significantly higher level of beige adipocyte markers (e.g. Pgc1a and Ucp1). |

For sentences that met the criteria for extracting cell–gene relationship pairs based on rules, we utilized the tagger and parse components of the SciBERT model to parse the syntactic structure of the sentences and generate a syntactic dependency tree (Fig. 3). This tree effectively illustrates the relationships between tokens. We extracted cell‒gene relationship pairs that were located within the same subtree. A subtree was defined as a sequence that included the token and all its syntactic descendants. Additionally, it is worth mentioning that sentence structures that conformed to the pattern “cell name (gene name)” were directly selected for the extraction of cell–gene pairs.

**Fig. 3.**
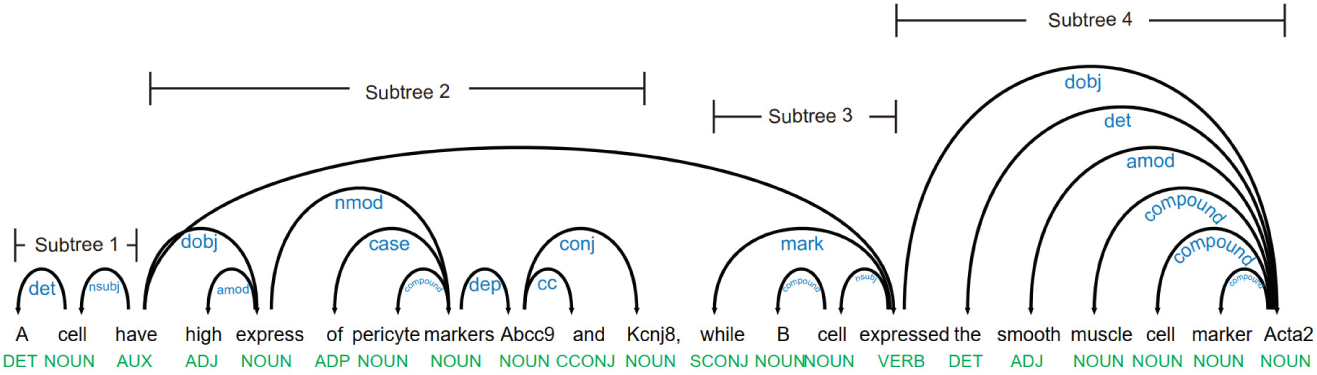
An example of a syntactic dependency tree.

### 3.5 Statistics

All the statistical analyses were performed in R (version 4.1). The performance of the marker-related sentence classification model was evaluated using the precision, recall, and F1 score of the predicted entity tag compared to the ground truth labels.

## 4. Results

### 4.1 Identification of gene and cell entities using MarkerGeneBERT

Pretrained NER models for entity extraction have proven to be effective in various research fields. MarkerGeneBERT integrates three pretrained NER models based on diverse biomedical corpora. Additionally, we incorporated cell names curated from the Cell Ontology database for exact string matching. Given the standardized gene names, the MarkerGeneBERT utilized only gene symbol IDs exclusively sourced from the GTF file in Cell Ranger for accurate gene entity recognition. Further details can be found in the Methods section.

As described in the Methods section, 27323 sentences labeled with cell and gene names were manually annotated by our team and originally designed for the marker-related sentence classification model. These were also used for validation of cell and gene entity identification performance of “en_ner_bionlp13cg_md”, “en_ner_craft_md”, “en_ner_jnlpba_md” and MarkerGeneBERT. Compared to the three pretrained NER models used alone, the MarkerGeneBERT achieved higher precision and recall in the extraction of cell and gene names (Table 4). For gene name identification, the MarkerGeneBERT achieved an F1 score of 87% (precision: 89%, recall: 99%), which is 20% greater than that of the second-best model. Regarding cell name identification, the MarkerGeneBERT obtained an F1 score of 92% (precision: 86%, recall: 98%), surpassing the second-best model by 8%. demonstrating the best trade-off between precision and recall.

**Table 4.** Performance of various NER models on the manually annotated dataset.

| NER model | Gene entity |  |  | Cell type entity |  |  |
| --- | --- | --- | --- | --- | --- | --- |
|  | P (%) | R (%) | F1 (%) | P (%) | R (%) | F1 (%) |
| en_ner_bionlp13cg_md | 58 | 80 | 67 | 84 | 82 | 83 |
| en_ner_craft_md | 64 | 70 | 67 | 93 | 58 | 72 |
| en_ner_jnlpba_md | - | - | - | 90 | 79 | 84 |
| MarkerGeneBERT (ours) | 78 | 99 | 087 | 86 | 98 | 92 |

### 4.2 Cell–biomarker associative binary classification

We introduced a supervised marker-related text classification model to determine which sentences included not only cell entities and gene entities but also specific syntactic patterns indicating that a gene is a marker of a cell. More details about the model and training dataset construction process are available in the Methods section.

To evaluate the performance of the marker-related text classification model in distinguishing specific syntactic patterns indicating that genes are markers of cells, we partitioned the training dataset into 10 subsets, randomly selecting 9 subsets for model training and reserving one subset for validation. The evaluation results depicted in Fig. 4A demonstrated a mean average precision (mAP) of 0.876 (ranging from 0.84 to 0.91), a mean precision of 0.844 (ranging from 0.8 to 0.9), and a mean recall of 0.734 (ranging from 0.56 to 0.78).

**Fig. 4.**
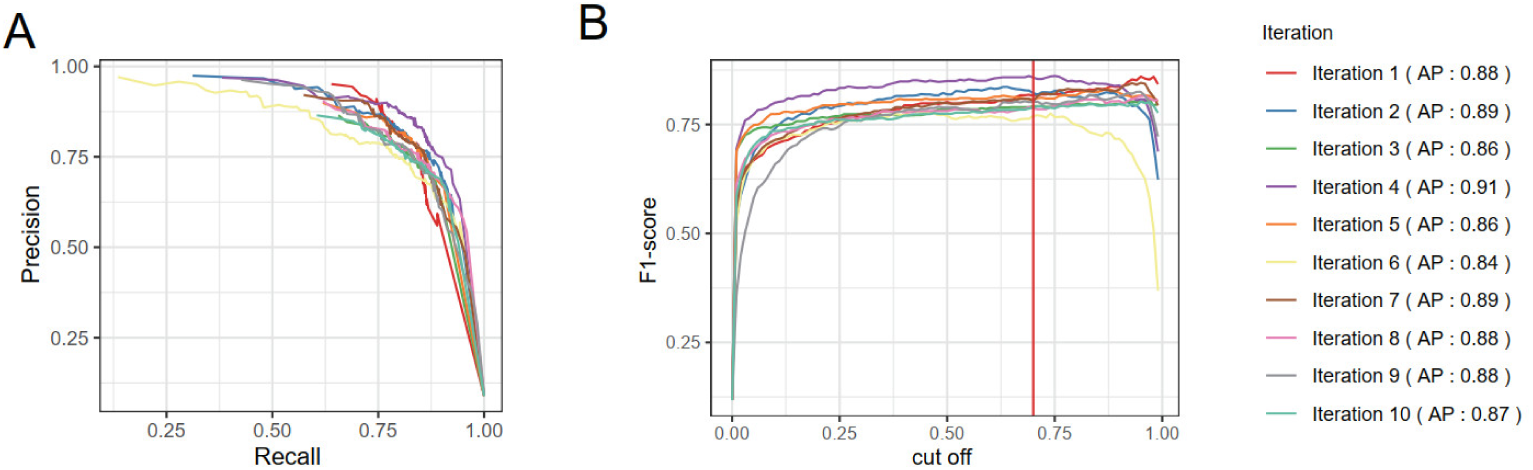
Evaluation of the marker-related text classification model. **A** Precision‒recall curve for the training model on the validation set at various iterations. **B** F1-scores for the training cohort in the validation cohort for different cutoff values. The mean F1-score is greatest at the vertical red line.

After processing by the model, each sentence could obtain a predicted probability value. A sentence was classified as a marker-related sentence if the predicted probability value was greater than the threshold, so the threshold setting was very important for the performance of our model. We calculated the F1 score for different thresholds, as illustrated in Fig. 4B, and the fitting threshold was 0.7. Under these threshold settings, the F1 score achieved optimal performance across different validation sets.

For the remaining marker-related sentences whose predicted probability was greater than 0.7, we employed syntactic structure-based analysis within each sentence to identify and extract reliable cell-marker relationship pairs. The extraction criteria are described in detail in the Methods section. In addition, we employed an appropriate NER model, as shown in Table 2, to assess the species, organs, and disease information in each study. Further details are provided in the Methods section.

### 4.3 Statistics of the NLP system extraction results

We employed MarkerGeneBERT to extract approximately 4000 cell types and approximately 20000 genes from 3987 literature sources (Supplemental Table 1). Compared to existing databases manually curated by domain experts over the years, our model achieved competitive retrieval results (Table 5). The maximum memory of our system, which included all the scripts and models, was 21 GB, and the parsing and entity extraction of one paper could be quickly completed in 7 minutes.

**Table 5.** Comparison of inclusion results across different databases.

| Database | Literature | Source | Type | Date | Species | Tissue | Disease | Cell | Marker |
| --- | --- | --- | --- | --- | --- | --- | --- | --- | --- |
| CellMarker2.0 | scRNA-seq (1945) | Full text | manual | 2022 | Human | 828 | 374 | 3149 | 27122 |
|  | & Experiment (4040) |  | curated |  | & Mouse |  | (cancer |  |  |
|  | & Review (354) |  |  |  |  |  | type) |  |  |
| Panglao DB | scRNA-seq (1054) | Full text | manual | 2020.3 | Human | 29 | — | 178 | 8286 |
|  |  |  | curated |  | & Mouse |  |  |  |  |
| scALE | 26 million | main text | NLP | 2021 | Human | Pan | — | 556 | — |
|  | biomedical |  |  |  | & Mouse | tissue |  |  |  |
|  | documents |  |  |  |  |  |  |  |  |
| MarkerGeneBERT | scRNA-seq (3987) | main text | NLP | 2023.6 | Human | 744 | 737 | 3565 | 16875 |
|  |  |  |  |  | & Mouse |  |  |  |  |

### 4.4 Concordance between MarkerGeneBERT and manually curated databases

To validate the accuracy of the cell entity, gene entity, cell-marker pair, species, tissue, and disease information detection system, we compared the results with those of CellMarker2.0, which is widely utilized as the gold standard for manual curation. Given that our methodology primarily extracted gene markers from the main text, we specifically compared the results from 1027 articles that were in both CellMarker2.0 and our database. Other articles were excluded for reasons such as unavailability for download or because the markers were sourced from supplemental materials; further details can be found in Supplemental Fig. 1.

#### 4.4.1 The MarkerGeneBERT identifies most cell and gene entities recorded in databases

In the common 1027 studies, the CellMarker2.0 manual curated a total of 4646 cell types with 12874 marker genes, while the main text parts covered 3185 cell types and 8683 marker genes; approximately 84% of the valuable information was derived from the main text (Supplemental Fig. 2). MarkerGeneBERT identified 90.8% of the marker gene entities (7890/8683) and 92.7% of the cell type entities (2954/3185) in these common studies (Fig. 5A).

**Fig. 5.**
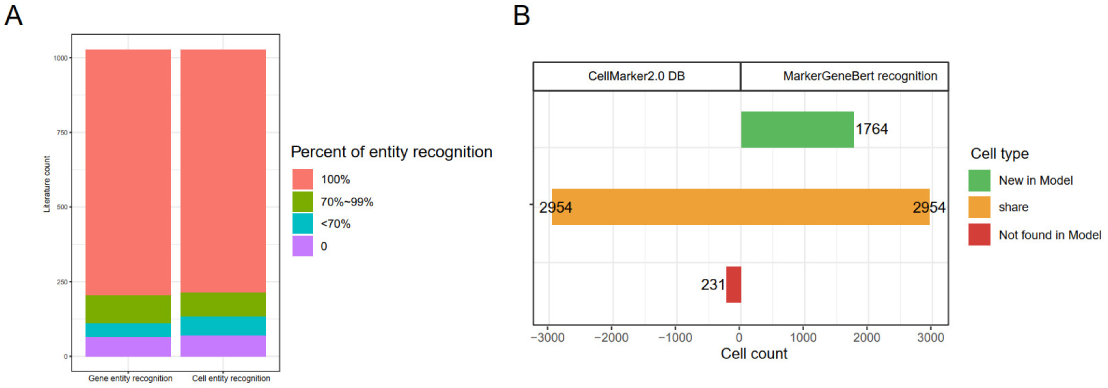
Stat in cells and gene entities recognized by MarkerGeneBERT. **A** Proportion of genes and cell types curated in the CellMarker2.0 database identified by MarkerGeneBERT. **B** Cell types recognized by MarkerGeneBERT

Through a systematic comparison of the results extracted from each literature source with those of CellMarker2.0, MarkerGeneBERT revealed an additional 1764 cell types associated with the marker genes (Fig. 5B). Among the 1764 newly identified cell types, 1344 were initially excluded by CellMarker2.0 in the corresponding literature; however, these were reported in other studies of the same tissue.

Notably, there were 89 cell types not cataloged in CellMarker2.0; these were mainly tissue-specific cell types, but the frequencies of these cells, such as enteric mesothelial fibroblasts from the intestine and retinal progenitor cells from the ocular tissue, were low. Additionally, 302 cell types were detected with CellMarker2.0 but not with corresponding tissues. We categorized these 89 newly recorded cell types and 302 reported cell types according to their tissue information (Fig. 6). These cell types primarily represent functional cells distributed across different tissues; for instance, in the literature related to human gastric tissue, cancer-associated fibroblasts (CAFs), as central components of the tumor microenvironment in primary and metastatic tumors, profoundly influence the behavior of cancer cells and are involved in cancer progression through extensive interactions with cancer cells and other stromal cells[25]. Our method can be used to directly record CAFs in both cancer and gastric tissues. The detailed cell marker information is available in Supplemental Table 2.

**Fig. 6.**
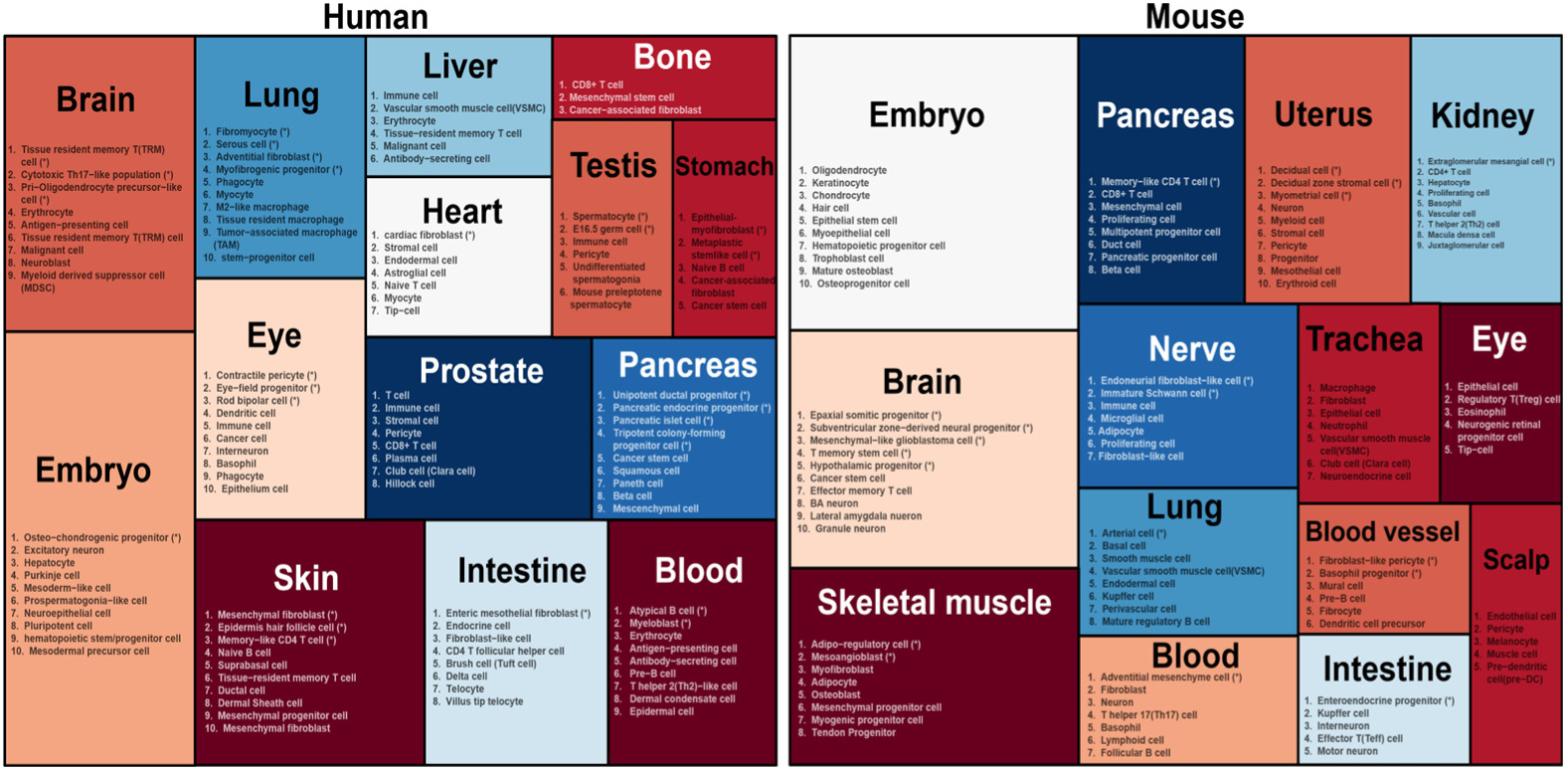
MarkerGeneBERT identified novel cell types in specific tissues. Cell types marked with a suffix * were those not documented in any tissue type within the CellMarker2.0 database.

#### 4.4.2 High consistency of the marker gene list between the MarkerGeneBERT and the database

For each study, we assessed the consistency of the cell marker genes identified between CellMarker2.0 and MarkerGeneBERT. As illustrated in Fig. 7, approximately 47% of the cell types and their corresponding marker gene pairs were the same in the CellMarker2.0 database and MarkerGeneBERT. Additionally, for approximately 23% of the cell types, the marker genes extracted by MarkerGeneBERT were present in CellMarker2.0, and they accounted for 87% of the corresponding marker genes recorded in CellMarker2.0. The reason for the extraction results falling short of 100% was primarily due to certain cell types that record multiple marker genes within a single document, and it was possible that MarkerGeneBERT may have filtered out some marker genes based on preset conditions (Supplemental Fig. 3). And still, the majority of such cell markers also showed a high level of precision, often reaching 100%. Overall, MarkerGeneBERT exhibited a high percentage of true positives, and there was a high level of consistency between the results extracted from the MarkerGeneBERT and CellMarker2.0 databases.

**Fig. 7.**
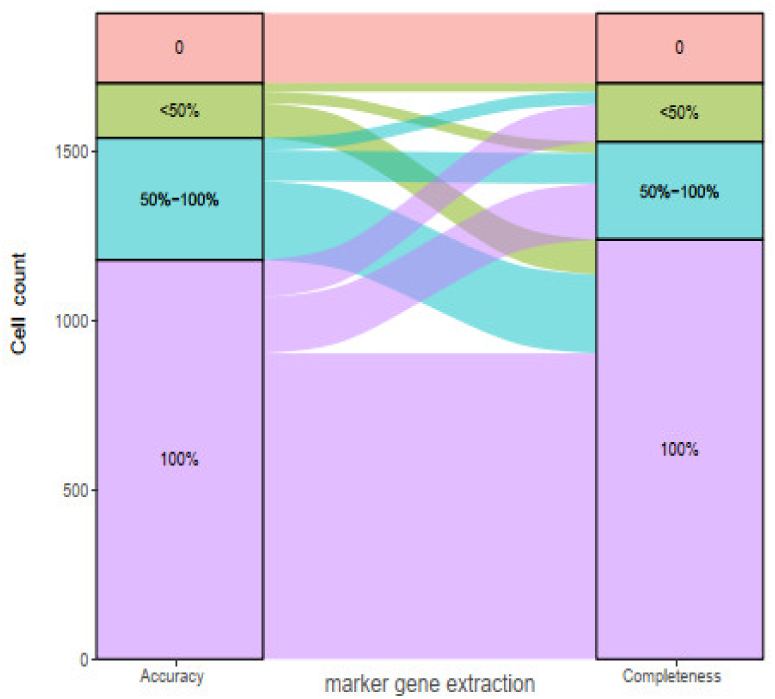
The completeness and accuracy of marker extraction for each cell in every st Accuracy: For a cell, the proportion of the intersection of the number of markers collected in the Cellmarker2.0 database and the number of markers extracted by the NLP-based model in a specific literature to the number of markers extracted by the model Completeness: For a cell, the proportion of the intersection of the number of markers collected in the CellMarker2.0 database and the number of markers extracted by the model in a specific literature to the number of markers collected in the Cellmarker2.0 database

Additionally, approximately 13% of the cells and their marker genes reported in CellMarker2.0 were 100% of those found by MarkerGeneBERT, and on average, MarkerGeneBERT obtained 25% more marker genes that were not recorded by CellMarker2.0. We traced back some newly discovered marker genes in the original text and found that CellMarker2.0 may ignore marker genes inconsistent with the main research themes of the paper or that only the first half of the information was extracted, while the following half was ignored.

#### 4.4.3 Consistency of species, tissue, and disease

We compared the consistency of species, tissue and disease information extracted from 1540 studies between the NLP system and CellMarker2.0. Overall, the consistency rates were 75% for species information, 77% for tissue information, and 66% for disease information.

The primary reason for the lower-than-expected consistency stemmed from our emphasis on organizing and analyzing information extracted from the full texts of the specific studies, summarizing the main species, tissues, and disease types studied. In contrast, the CellMarker2.0 database uses literature IDs as indices to trace cell markers referenced from other literature sources, capturing the associated species, tissue, and disease information from both reference and specific literature. Consequently, there is variance in the information recorded by these two methods in the same study.

### 4.5 Increased cell type annotation efficiency through multi-marker annotation strategies

MarkerGeneBERT collected 166 brain cell types from approximately 190 studies, including some cell types not previously cataloged in CellMarker2.0, such as tissue-resident memory T cells, neuroblasts, and myeloid-derived suppressor cells (Supplemental Table 3). We utilized these 166 brain cell types and their compiled marker gene lists on published posterior hippocampus single-cell RNA data for cell type annotation by using scCATCH, which is a cell type annotation tool based on preset marker gene list. As illustrated in Fig. 8A, the cell type annotations obtained directly by scCATCH were almost the same as those in the original paper labels[26]. Notably, among the top 5 differentially expressed genes(DEGs) identified by scCATCH for cell type annotation, seven were newly discovered in our database and not recorded in CellMarker2.0 (Fig. 8B). This indicates that while many cell types possess representative marker genes, such as the CD3 marker for immune cells, which is mentioned and used in numerous articles, a more comprehensive list of marker genes can enhance the annotation efficiency of automated cell type annotation methods or tools.

**Fig. 8.**
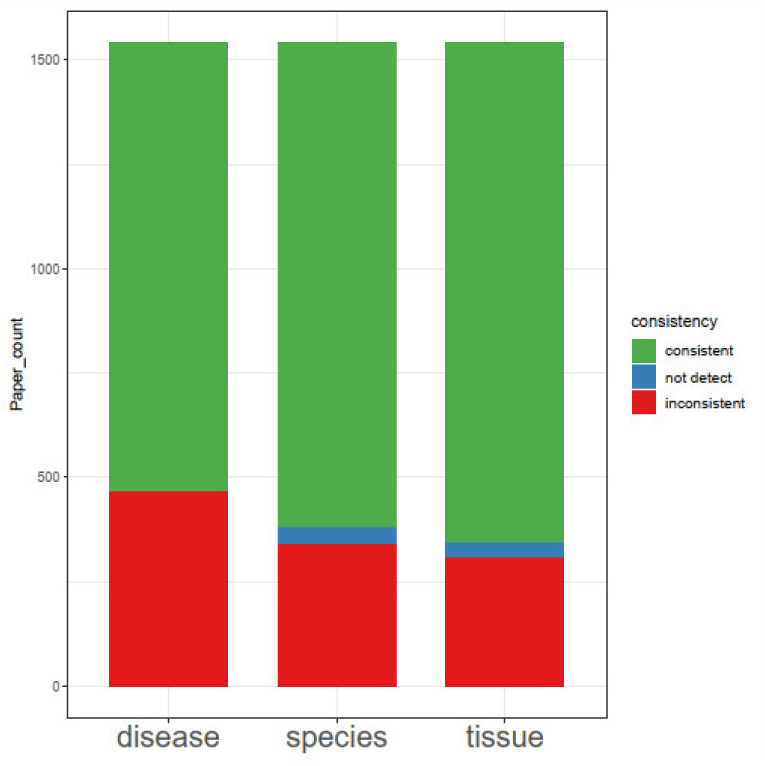
Consistency between the species, tissue, and disease types recorded in the CellMarker2.0 and those inferred by the MarkerGeneBERT.

**Fig. 9.**
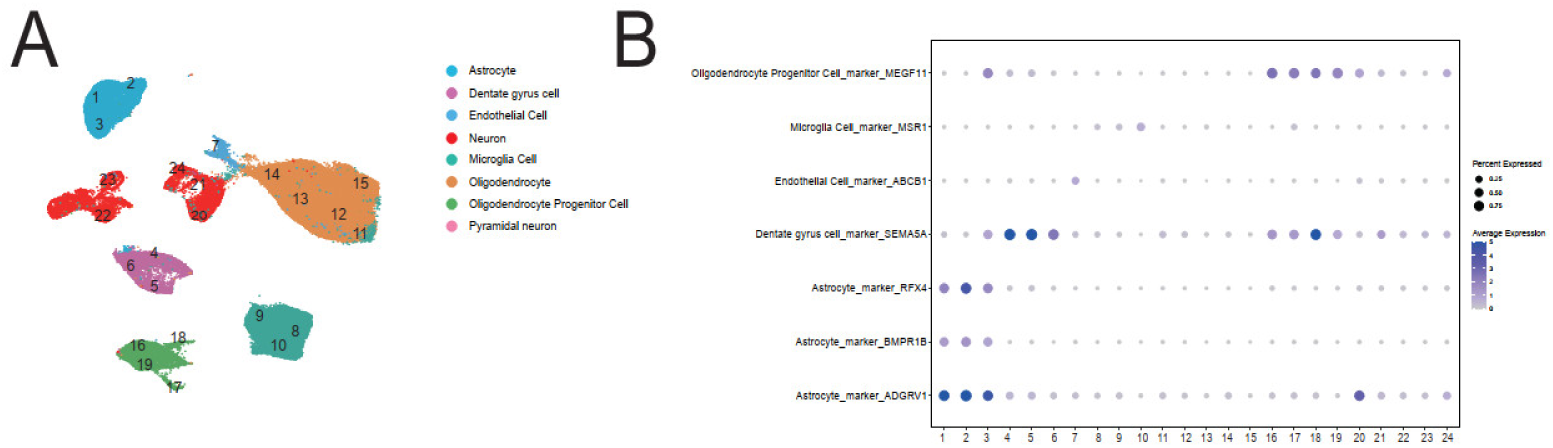
Consistency between the cell type annotation results of single-cell sequencing in hippopotamus tissue and the original annotation results[26] **A** Cell type annotation results of single-cell sequencing in hippopotamus tissue. **B** The seven top DEGs used for scCATCH cell type annotation were newly extracted by MarkerGeneBERT.

## 5. Discussion

In the coming years, single-cell sequencing technology is expected to be applied to a wider range of species and tissue types. This necessitates that researchers possess effective capabilities for annotating and analyzing such data. Although there are several manually curated marker gene databases and methods for constructing corresponding databases, manual curation remains prohibitively time-consuming and introduces potential biases, particularly when confronted with complex cell type annotations.

The developed NLP models for extracting entity relationships from text have been extensively applied across diverse domains. Doddahonnaiah *et al*. utilized the co-occurring theory to calculate the co-occurring frequencies of 500 pre-selected cell names and common genes in biomedical documents, thereby inferring cell marker genes[15]. However, as discussed in their work and as mentioned in a related study[13], the co-occurring theory itself has several limitations, as a pair of entities with low co-occurring frequencies can be reliable but may go undetected. Additionally, the pre-selection of cell names significantly constrains the scalability of the method. In contrast to co-occurrence methods, we optimized a dependency analysis method based on NLP models to capture all mentioned cell names within the text and extract cell–gene pairs that are related in grammatical structure. Additionally, utilizing artificially annotated marker-related sentences created by our team, we developed a marker-related text classification model to ensure that sentences containing cell–gene pairs are inherently marker related, thereby filtering out cell–gene pairs merely co-occurring within a sentence.

This study has several limitations. First, we utilized only the main text of the literature for extracting cell marker genes; however, in some related studies, certain cell markers are presented in the figures or Supplemental materials. Additionally, there was no standard tissue nomenclature, such as the use of “ascites” in single-cell studies, where it is frequently mentioned but not considered a tissue. We currently provide a pattern for two candidate tissues to meet the automated extraction needs for tissue information as much as possible. Unlike the recognition of cell and gene entities, correct tissue information typically relies on a comprehensive understanding of the entire text. Although we attempted to extract tissue information separately from the titles, abstracts, results, and methods sections, we relied on the frequency of different tissues to select the appropriate true tissue; in some cases, the tissues extracted were still erroneous, so the accuracy of extracting organizational information was not particularly high. In addition, to balance computational resources and time, we analyzed only human and mouse scRNA-seq literatures, as the gene names of different species are not completely the same, existing data preparation for humans and mouse cannot be directly applied to the extraction of new species. If there is a need to extract cell marker genes from the literature from other species, it is necessary to organize and construct a new gene dictionary. Finally, the most apparent advantage of, NLP system is its ability to swiftly extract sentences containing key terms (cell, and gene) and determine their associative relationships, there is no unified name or classification system for cells, and the cell names is largely influenced by personal writing habits, such as T cells, t cell, B cells, CD4+ T cells, cd4 t cells, and CD8+ T cells. For better construction of cell marker databases, classifying different cell names according to the true cell type remains an enormous challenge for both human manual and NLP automated methods.

During the comparison of the cell markers MarkerGeneBERT and Cellmarker2.0, 802 cell types were detected in Cellmarker2.0 and MarkerGeneBERT, but their expression was classified as non-marker-related due to the significantly lower probability of being predicted by the model (Supplemental Fig. 4). In addition, the texts of 242 other cell types met the model threshold. However, the complex grammatical structures of these cells currently challenge our methods for identifying cell-marker pairs within them. Considering the above issues, our goal in the future is to incorporate a more diverse training dataset for marker-related text classification model training to further accommodate the screening of diverse marker-related texts. Moreover, we plan to introduce a structured binary graph transformer (SBGT) model similar to the model constructed and trained by Gu et al. [13] to further improve the accuracy of entity relationship determination.

## 6. Conclusion

We developed an NLP-based text mining system named MarkerGeneBERT to identify cells and marker genes from both the text and tables in the literature sourced from PubMed and PubMed Central. Using artificially annotated marker-related sentences, we constructed a supervised text classification model to initially screen out texts containing both gene and cell names; then, cell and marker genes were extracted from those texts according to marker-related patterns. Our cell marker gene identification pipeline achieved 75% precision and 76% recall when compared with CellMarker2.0, demonstrating the success and state-of-the-art of cell marker gene extraction by NLP text mining.

## 7. Declarations

## Ethics approval and consent to participate

Not applicable

## Consent for publication

Not applicable

## Availability of data and materials

All data generated or analysed during this study are included in this published article and its supplemental information files. Transcriptome profiling of hippopotamus are available at GSE160189 (https://www.ncbi.nlm.nih.gov/geo/query/acc.cgi?acc=GSE160189)

## Competing interests

The authors declare that they have no competing interests.

## Funding

Not applicable

## Authors’ contributions

Conceptualization, XL Zhang, Peng Cheng, Yan Peng and YZ Li; methodology, Peng Cheng and XL Zhang; formal analysis, Peng Cheng; investigation, XL Zhang; data curation, Peng Yan and YZ Li; writing—Sheng Chen and Peng Cheng; writing—review and editing, Peng Yan and YZ Li; funding acquisition, Yan Peng and YZ Li. All authors read and approved the final manuscript.

## Acknowledgements

Not applicable

## Notes

### Competing Interest Statement

The authors have declared no competing interest.

## References

1. Jovic D, Liang X, Zeng H, Lin L, Xu F, Luo Y: Single-cell RNA sequencing technologies and applications: A brief overview. Clin Transl Med 2022, 12(3):e694.

2. Zhang AW, O’Flanagan C, Chavez EA, Lim JLP, Ceglia N, McPherson A, Wiens M, Walters P, Chan T, Hewitson B et al: Probabilistic cell-type assignment of single-cell RNA-seq for tumor microenvironment profiling. Nat Methods 2019, 16(10):1007–1015.

3. Shao X, Liao J, Lu X, Xue R, Ai N, Fan X: scCATCH: Automatic Annotation on Cell Types of Clusters from Single-Cell RNA Sequencing Data. iScience 2020, 23(3):100882.

4. Aran D, Looney AP, Liu L, Wu E, Fong V, Hsu A, Chak S, Naikawadi RP, Wolters PJ, Abate AR et al: Reference-based analysis of lung single-cell sequencing reveals a transitional profibrotic macrophage. Nat Immunol 2019, 20(2):163–172.

5. Pliner HA, Shendure J, Trapnell C: Supervised classification enables rapid annotation of cell atlases. Nat Methods 2019, 16(10):983–986.

6. Cao ZJ, Wei L, Lu S, Yang DC, Gao G: Searching large-scale scRNA-seq databases via unbiased cell embedding with Cell BLAST. Nat Commun 2020, 11(1):3458.

7. Hu C, Li T, Xu Y, Zhang X, Li F, Bai J, Chen J, Jiang W, Yang K, Ou Q et al: CellMarker 2.0: an updated database of manually curated cell markers in human/mouse and web tools based on scRNA-seq data. Nucleic Acids Res 2023, 51(D1):D870–D876.

8. Franzen O, Gan LM, Bjorkegren JLM: PanglaoDB: a web server for exploration of mouse and human single-cell RNA sequencing data. Database (Oxford) 2019, 2019.

9. Meng FL, Huang XL, Qin WY, Liu KB, Wang Y, Li M, Ren YH, Li YZ, Sun YM: singleCellBase: a high-quality manually curated database of cell markers for single cell annotation across multiple species. Biomark Res 2023, 11(1):83.

10. Jin J, Lu P, Xu Y, Tao J, Li Z, Wang S, Yu S, Wang C, Xie X, Gao J et al: PCMDB: a curated and comprehensive resource of plant cell markers. Nucleic Acids Res 2022, 50(D1):D1448–D1455.

11. Yuan H, Yan M, Zhang G, Liu W, Deng C, Liao G, Xu L, Luo T, Yan H, Long Z et al: CancerSEA: a cancer single-cell state atlas. Nucleic Acids Res 2019, 47(D1):D900–D908.

12. Shetty P, Rajan AC, Kuenneth C, Gupta S, Panchumarti LP, Holm L, Zhang C, Ramprasad R: A general-purpose material property data extraction pipeline from large polymer corpora using natural language processing. NPJ Comput Mater 2023, 9(1):52.

13. Gu W, Yang X, Yang M, Han K, Pan W, Zhu Z: MarkerGenie: an NLP-enabled text-mining system for biomedical entity relation extraction. Bioinform Adv 2022, 2(1):vbac035.

14. Naseri H, Kafi K, Skamene S, Tolba M, Faye MD, Ramia P, Khriguian J, Kildea J: Development of a generalizable natural language processing pipeline to extract physician-reported pain from clinical reports: Generated using publicly-available datasets and tested on institutional clinical reports for cancer patients with bone metastases. J Biomed Inform 2021, 120:103864.

15. Doddahonnaiah D, Lenehan PJ, Hughes TK, Zemmour D, Garcia-Rivera E, Venkatakrishnan AJ, Chilaka R, Khare A, Kasaraneni A, Garg A et al: A Literature-Derived Knowledge Graph Augments the Interpretation of Single Cell RNA-seq Datasets. Genes (Basel) 2021, 12(6).

16. Bada M, Eckert M, Evans D, Garcia K, Shipley K, Sitnikov D, Baumgartner WA, Jr., Cohen KB, Verspoor K, Blake JA et al: Concept annotation in the CRAFT corpus. BMC Bioinformatics 2012, 13:161.

17. Collier N, Kim J-D: Introduction to the bio-entity recognition task at JNLPBA. In: Proceedings of the International Joint Workshop on Natural Language Processing in Biomedicine and its Applications (NLPBA/BioNLP): 2004; 2004: 73–78.

18. Pyysalo S, Ohta T, Rak R, Rowley A, Chun HW, Jung SJ, Choi SP, Tsujii J, Ananiadou S: Overview of the Cancer Genetics and Pathway Curation tasks of BioNLP Shared Task 2013. BMC Bioinformatics 2015, 16 Suppl 10(Suppl 10):S2.

19. Kovalchik SJRpv: Download content from NCBI databases. 2014, 4(0):2021.

20. Fantini D, Fantini MD: Package ‘easy PubMed’. In.: CRAN; 2017.

21. Honnibal M, Montani IJTa: spaCy 2: Natural language understanding with Bloom embeddings, convolutional neural networks and incremental parsing. 2017, 7(1):411–420.

22. Neumann M, King D, Beltagy I, Ammar WJapa: ScispaCy: fast and robust models for biomedical natural language processing. 2019.

23. Beltagy I, Lo K, Cohan AJapa: SciBERT: A pretrained language model for scientific text. 2019.

24. Diehl AD, Meehan TF, Bradford YM, Brush MH, Dahdul WM, Dougall DS, He Y, Osumi-Sutherland D, Ruttenberg A, Sarntivijai S et al: The Cell Ontology 2016: enhanced content, modularization, and ontology interoperability. J Biomed Semantics 2016, 7(1):44.

25. Yang D, Liu J, Qian H, Zhuang Q: Cancer-associated fibroblasts: from basic science to anticancer therapy. Exp Mol Med 2023, 55(7):1322–1332.

26. Ayhan F, Kulkarni A, Berto S, Sivaprakasam K, Douglas C, Lega BC, Konopka G: Resolving cellular and molecular diversity along the hippocampal anterior-to-posterior axis in humans. Neuron 2021, 109(13):2091–2105 e2096.

