## Supplementary figures and images for "A natural language processing system for the efficient extraction of cell markers"

### Supplementarl Figure1

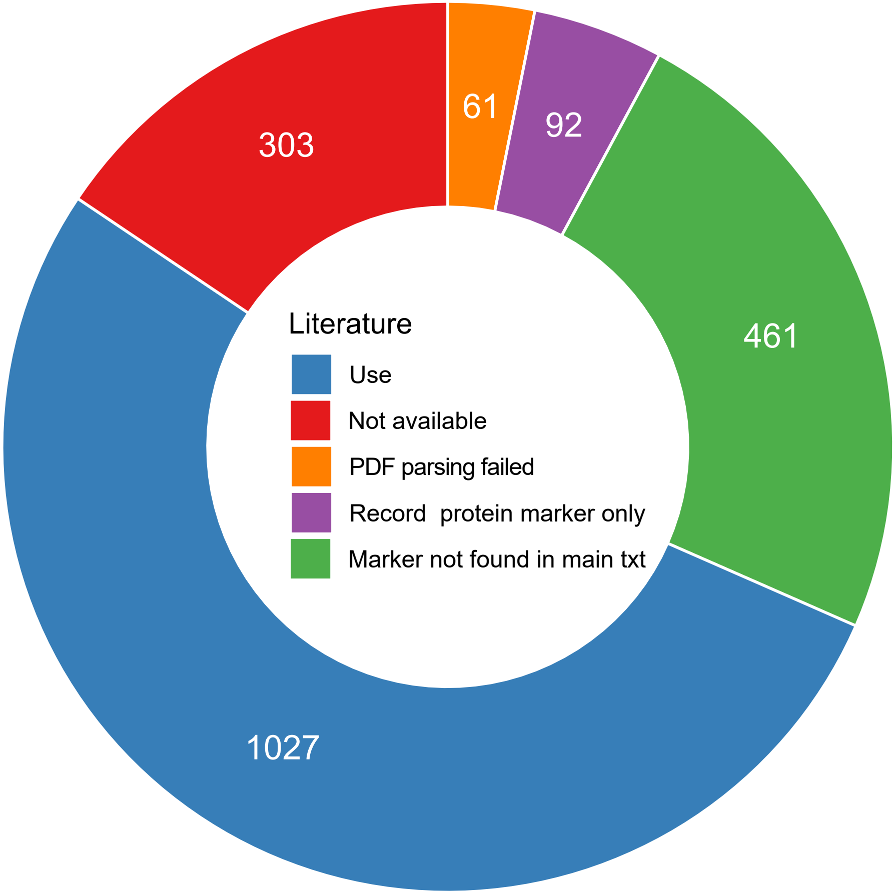

### Supplementarl Figure2

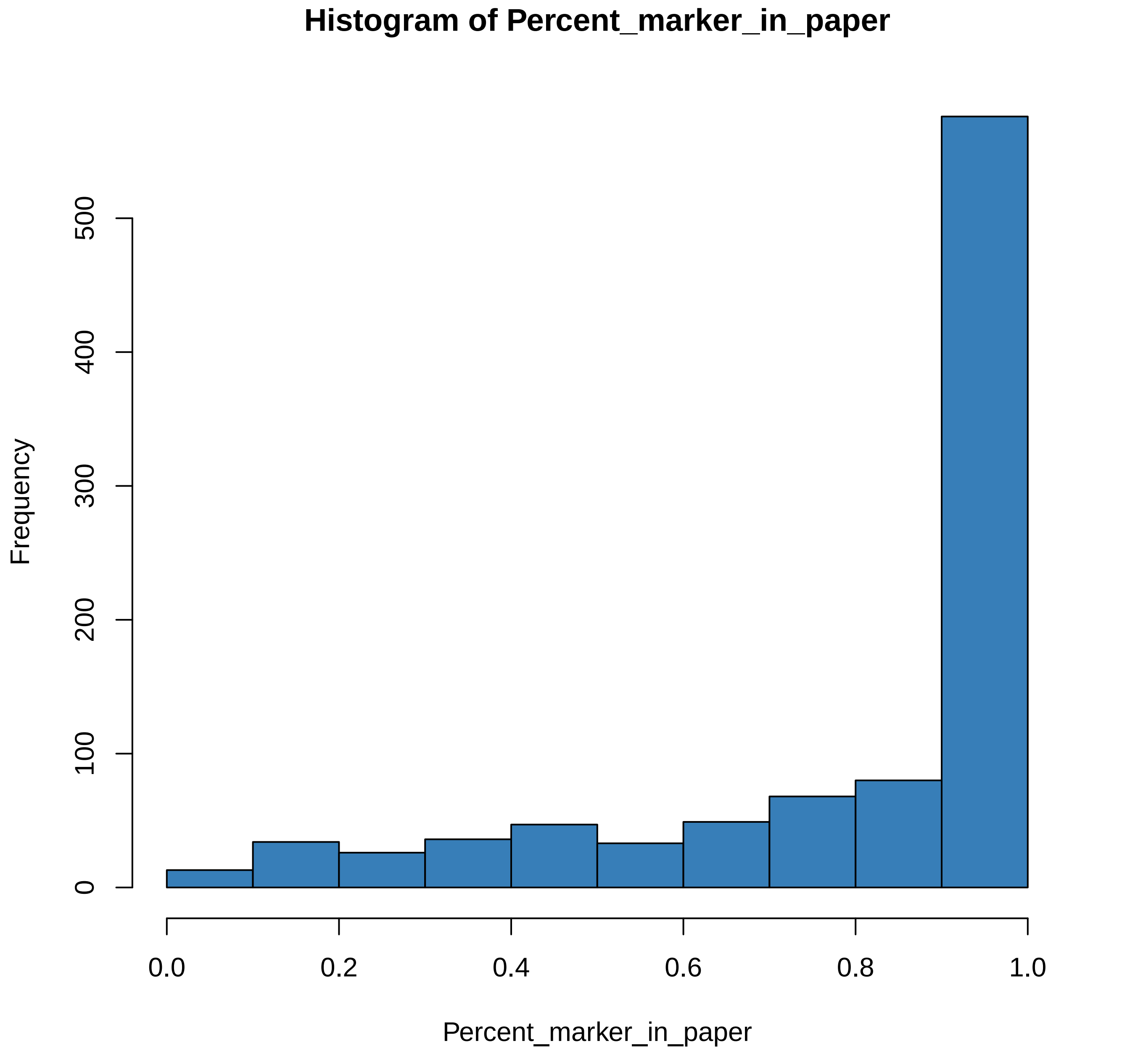

### Supplementarl Figure3

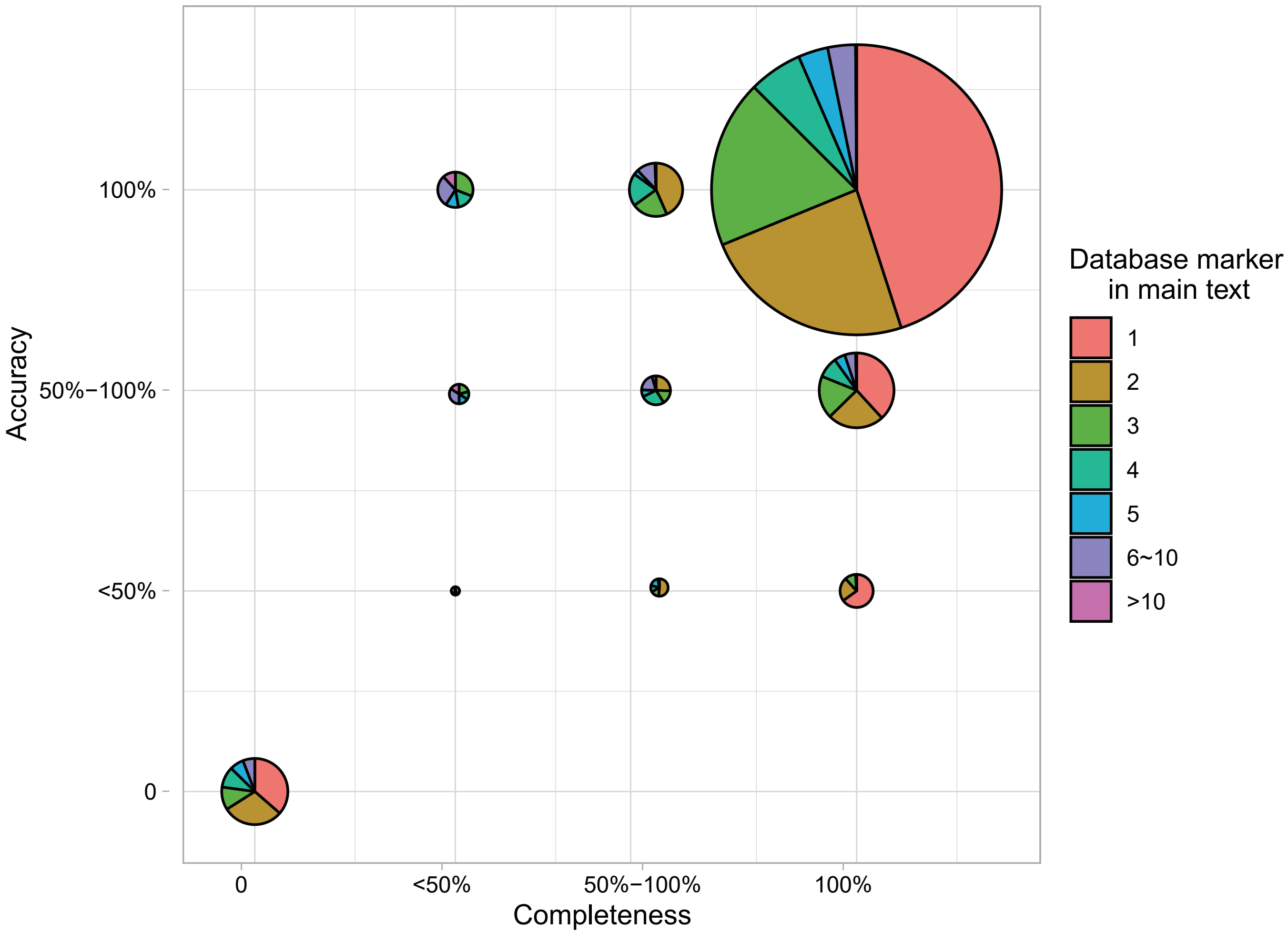

### Supplementarl Figure4

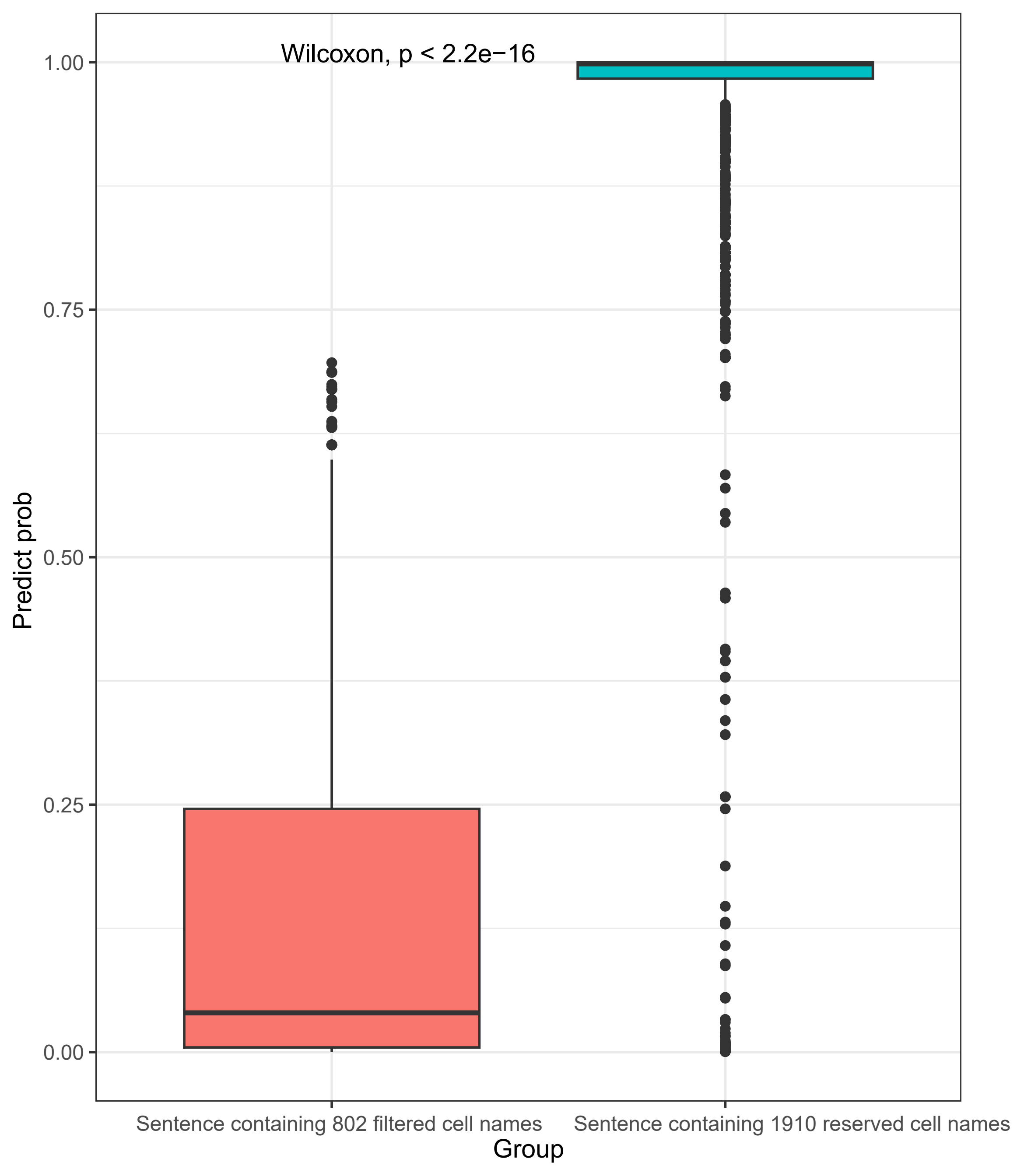
